# Expanding the repertoire of suicidal germination stimulants for control of parasitic weeds

**DOI:** 10.64898/2026.04.13.718129

**Authors:** Gracia D. Mave, Thommas M Musyoka, Sylvia M. Mutinda, Francisca Mutindi, Willy Kibet, Esther Mary Munyoki Toili, Samuel Muiruri, Justus Onguso, Jaindra Tripathi, Leena Tripathi, Steven Runo

## Abstract

Witchweeds (*Striga spp*.) are parasitic plants that severely constrain cereal production across sub-Saharan Africa, threatening food security for millions of people (Runo and Kuria, 2018). *Striga* infection begins when dormant seeds germinate in response to host-derived biomolecules, primarily strigolactones which plants emit to regulate shoot branching and to communicate with beneficial microbes. This obligate dependence on host signals can be exploited for *Striga* control through suicidal germination, whereby strigolactone-like compounds induce parasite germination in the absence of a host. Although this strategy proved highly effective during *Striga* eradication efforts in the United States using ethylene gas as a *Striga* germination inducer (Eplee, 1975; Iverson et al., 2011), its deployment in Africa has been limited by capacity to synthesize cost-effective strigolactone-like *Striga* germination inducers. Here, we show that structure-guided *in silico* screening of chemical libraries using AlphaFold2-modeled receptor–ligand interactions improve the efficiency and likelihood of identifying previously unknown strigolactone analogs. Using this approach, we identify a structurally simple synthetic lactone scaffold that induces *Striga* germination at nanomolar concentrations. These results present new avenues for the development of strigolactone analogs and support revisiting suicidal germination as a practical *Striga* control strategy in Africa.

## Introduction

We propose that discovering new, simplified strigolactone analogs can be developed building on previous studies (4) that demonstrated that SL-based *Striga* germination requires only an aliphatic R group linked to a rigid C-ring and a D-ring lactone via an enol ether bridge, rather than the full ABCD tetracyclic scaffold. By screening already available vast small molecule libraries, compounds with SL-like features can be identified and designed. Virtual *in silico* screening using AlphaFold2 models of *Striga* germination receptors can then be performed: first to evaluate whether they recapitulate known strigolactone–receptor interactions, then to assess ligand stability through molecular dynamics simulations, prior to experimental validation. This strategy expands the accessible chemical search space and enhances the likelihood of identifying biologically active molecules.

## Results and Discussion

To test the viability of a structure-guided approach for identifying new strigolactone analogs, we performed a ligand-similarity search using Cartblanche22, a free database of commercially available compounds for virtual screening (5), with a GR24 scaffold lacking the B ring as the query. This search yielded 3-methyl-5-{[(3E)-2-oxo-5-phenyloxolan-3-ylidene]methoxy}-2,5-dihydrofuran-2-one and seven structural isomers. These compounds were designated Synthetic Lactone Analogs (SLAs) 1–8 (Fig. 1a).

**Figure 1.**
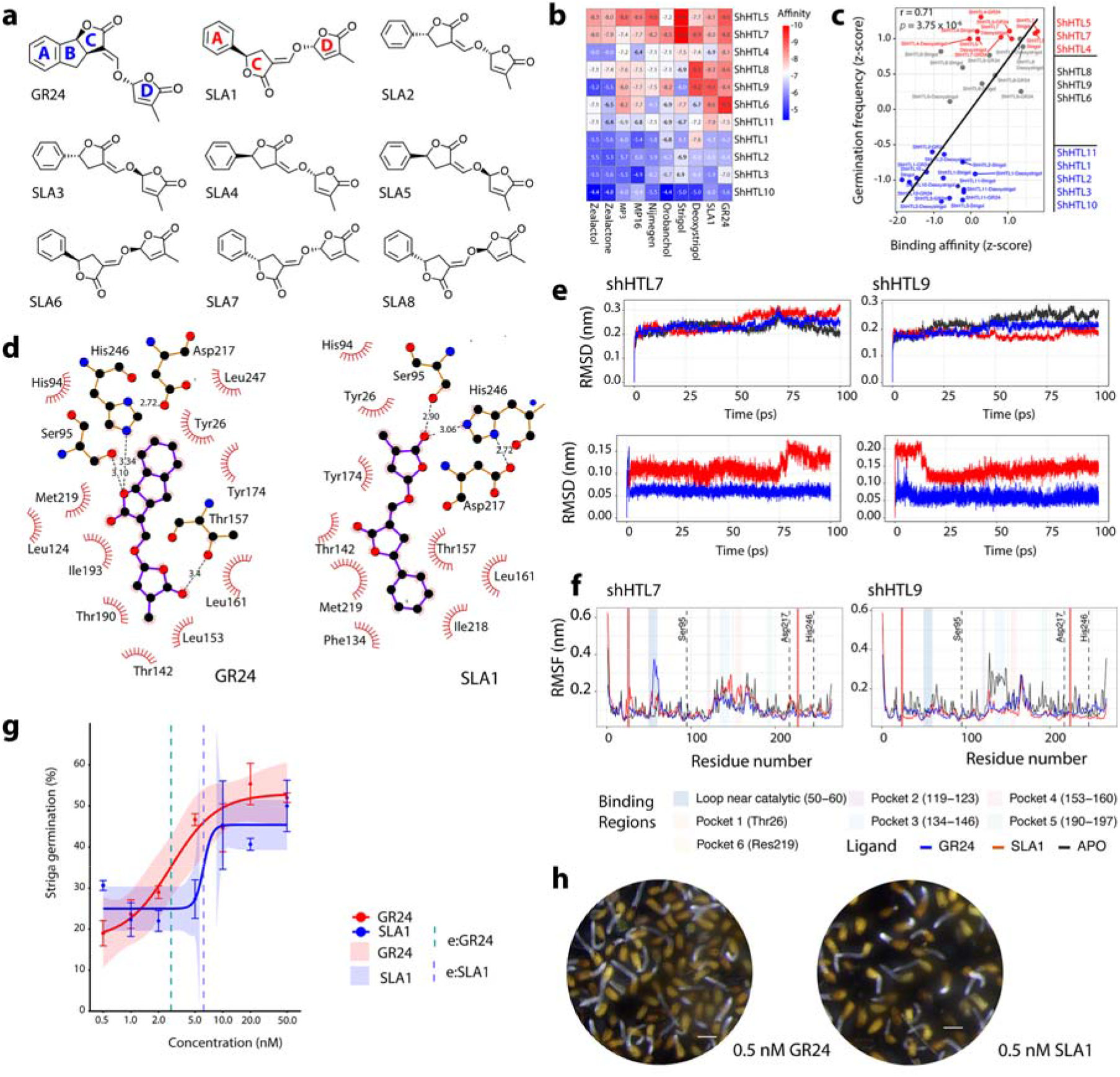
**(a)** Structures of GR24 and the synthetic lactone analogs (SLA1-8) identified in this study. Canonical strigolactones contain a tricyclic ABC scaffold linked to a D-ring lactone. SLA1 and its isomers retain the minimal A, C, and D ring architecture required for activity. **(b-d)** *In silico* docking analyses of strigolactones (SLs) and SL-like molecules with *Striga hermonthica* HYPERSENSITIVE TO LIGHT (ShHTL) receptors. **(b)** Heatmap of predicted docking affinities across ShHTL receptors. **(c)** Correlation between *Striga* germination stimulation and predicted binding affinities of SL-like molecules. **(d)** LIGPLOT schematics showing two-dimensional ligand– receptor interaction patterns for GR24 and SLA1, highlighting conserved interactions with Ser95 and His246, with Asp217 positioned to stabilize His246 within the catalytic triad. **(e-f)** Molecular dynamics (MD) simulations of ShHTL receptors in complex with GR24 or SLA1. **(e)** Root-mean-square deviation (RMSD) profiles of proteins and ligands. **(f)** Root-mean-square fluctuation (RMSF) of binding-site residues. **(g-h)** *Striga* germination assays. **(g)** Dose–response stimulation of *Striga* seed germination by SLA1 across serial dilutions, with GR24 as a positive control and (G nonlinear regression fits. (H) Representative images of germinated *Striga* seedlings treated with GR24 or SLA1 (scale = 0.5 cm).

To assess whether SLAs retained the structural features consistent with *Striga* germination stimulation, we performed molecular docking of SLA1 across all characterized *Striga hermonthica* HYPERSENSITIVE TO LIGHT (ShHTL) receptors and compared predicted binding affinities with previously reported germination responses of ShHTL-expressing *Arabidopsis* seeds exposed to various SLs and SL-like molecules (6). Predicted affinities (Fig. 1*b* and *c*) were strongly positively correlated with reported germination responses across ligands (r = 0.71, p = 3.75 × 10^−^□) (Fig. 1*b*). Canonical SLs exhibited the strongest predicted binding, including strigol (−10.0 kcal/mol on ShHTL7) and 5-deoxystrigol (−9.3 kcal/mol on ShHTL8 and ShHTL9). Among synthetic analogs, MP3, MP16, and Nijmegen also showed strong binding, particularly to ShHTL5 (−8.6 to −9.0 kcal/mol). In contrast, larger noncanonical SLs, such as zealactol, displayed weak affinities across receptors, except for ShHTL5, likely reflecting its larger ligand-binding pocket (7). Overall, these results support the predictive value of receptor–ligand binding affinities for germination activity. Based on its predicted binding, SLA1 is represents a plausible candidate, exhibiting affinities of −8.2 to −8.5 kcal/mol across ShHTL receptors, comparable to those of GR24.

Docking poses further supported the view that SLA1 retains SL-like structural features. SLA1 and GR24 occupied the same ligand-binding cavity and adopted closely related orientations within ShHTL receptors. On the high-sensitivity receptor ShHTL7, GR24 formed a hydrophobic pocket through its A, B, and C rings (Fig. 1*d*). SLA1 formed a similar core, with the A and C rings compensating for the absence of the B ring by occupying adjacent hydrophobic space (Fig. 1*d*). Importantly, both ligands retained hydrogen-bond interactions with Ser95 and His246 and contacted Asp217, confirming engagement of the complete Ser95–His246–Asp217 catalytic triad (8) (Fig. 1d and Movie S1 and S2).

Molecular dynamics (MD) simulations performed on ShHTL7 and ShHTL9 in complex with SLA1 or GR24 reinforced the prediction that SLA1 functions as an active SL-like analog (Movie S1 and S2). RMSD (root-mean-square deviation) analyses showed that the conformations of ligands, proteins, and complexes remained stable over time (Fig. 1e; Movie S4-S6). RMSF (root-mean-square fluctuation) showed minimal flexibility of the catalytic triad residues (Ser95, Asp217, His246; <0.08 nm) (Fig. 1*e*). And MM/GBSA (molecular mechanics/generalized Born surface area) calculations indicated comparable binding energies for SLA1 (−5436 ± 135 and −6059 ± 143 kcal/mol) and GR24 (−3889 ± 145 and −4424 ± 142 kcal/mol).

Consistent with computational predictions, *Striga* germination assays confirmed biological activity of SLA1 by germination induction in a dose-dependent manner, (Fig 1g and h) reaching an EC_50_ comparable to GR24 and reported activities of MP3 (5–20 nM) and Nijmegen-1 (10–50 nM) (9).

In sum, we show that structure-based computational screening encompassing ligand similarity searches, receptor docking, and molecular dynamics simulations was broadly predictive of *Striga* germination induction activity. Remaining challenges include modeling non-canonical agonists such as SPL7, whose strong germination activity extends beyond canonical Ser–His–Asp catalytic triad hydrolysis (10), developing high-throughput screening pipelines, and translating the validated *Striga* germination assays into effective field deployment.

## Materials and methods

Similarity-based searches for strigolactone-like small molecules were conducted in the Cartblanche database (5). Molecular docking was performed using AutoDock Vina (11) with *ShHTL* receptor structures predicted using ColabFold (12). Ligand structures were retrieved from PubChem; compounds not available were manually drawn in ChemDoodle (13). Molecular dynamics simulations were carried out using GROMACS 2024 (14). *Striga* germination assays were performed as previously described (15). See *SI Appendix* for details.

## Data and Software Availability

All scripts and raw data supporting this study are available at https://github.com/mutemibiochemistry/SLA1

## Acknowledgments

We acknowledge support from the African Union Commission through a PhD fellowship to G.D.M. High-performance computing resources were provided by the Centre for High Performance Computing (CHPC), South Africa.

## Supporting Information

### Extended Methods

#### Identification of synthetic lactone analogs

Similarity screening for strigolactone-like molecules was conducted in the CartBlanche22 database (https://cartblanche.docking.org/similarity/sw; accessed July 16, 2025) using a GR24 scaffold lacking the B ring as the query structure (1). The top-ranked compound, *3-methyl-5-{[(3E)-2-oxo-5-phenyloxolan-3-ylidene]methoxy}-2,5-dihydrofuran-2-one*, named Strigolactone Lactone 1 (SLA1) was selected for further analysis. Predicted binding affinities of this compound were evaluated and compared with those of canonical strigolactones and previously characterized strigolactone analogs across all *Striga hermonthica* HYPERSENSITIVE TO LIGHT (ShHTL) receptors. The resulting docking affinities were subsequently used in a correlation analysis with germination frequency data reported in (2).

#### Computational modeling and docking

The full repertoire of ShHTL receptors (2) was predicted using ColabFold (3) an open-source software platform developed in collaboration with Google that provides accessible protein structure prediction based on AlphaFold2 (4). For each receptor, the top-ranked structural model was selected for molecular docking. Structures of canonical strigolactones and established analogs were retrieved from PubChem (5). Ligands not available in PubChem were manually drawn using ChemDoodle (6), converted to three-dimensional conformations, and energy-minimized prior to docking. Molecular docking was performed using AutoDock Vina 1.2.0 (7) and predicted binding affinities were compared across all ShHTL receptors. Protein-ligand complexes were visualized using LigPlot+ v.2.3 (8) and The PyMOL Molecular Graphics System, Version 2.5 (9).

#### Molecular dynamics simulations and interaction energy estimations

To assess conformational stability of receptor–ligand complexes, 100-ns all-atom molecular dynamics (MD) simulations were performed for HTL7 and HTL9 bound to SLA1 or GR24 using GROMACS 2024.4 (10). Protein topologies were generated with the AMBER99SB-ILDN force field (11), while ligand parameters were prepared using the Antechamber Python Parser Interface (ACPYPE) (12). Receptor and ligand coordinates were combined under periodic boundary conditions, solvated in a triclinic water box with a 1.75-nm buffer, and energy-minimized using the steepest descent algorithm until the maximum force was <1000 kJ mol^−1^ nm^−1^.

Systems were equilibrated in two phases (100 ps each) to achieve stable temperature (300 K) and pressure (1 atm) using the modified Berendsen thermostat and the Parrinello–Rahman barostat (13). Production simulations were then conducted for 100 ns without position restraints, using a 2-fs integration time step, with an average computational cost of approximately 10,000 CPU hours per system. Trajectory analyses, including root-mean-square deviation (RMSD) and fluctuation (RMSF), were performed using standard GROMACS utilities, and plots were generated in R using ggplot2 (14). Binding free energies were estimated using the gmx-MMPBSA toolkit (v1.6.4) (15) to estimate the binding free energies (BFE) for the receptor-ligand complexes based on the energy variations between the bound and unbound states. For each system, 5,000 snapshot structures extracted from a converged 10-ns trajectory window were used to calculate binding free energies based on differences between bound and unbound states, following the protocol described in (16).

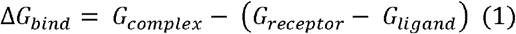

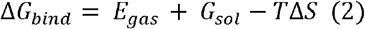

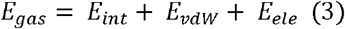

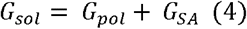

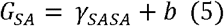

Where absolute free energies of the receptor-ligand complex, receptor and ligand are denoted by *G*_*complex*_, *G*_*receptor*_, *G*_*ligand*_ respectively. The binding free energy (Δ*G*_*bind*_) was decomposed to the various energetic contributions as indicated in eqn 2, 3, 4 and 5.

### *Striga* germination assays

*Striga hermonthica* seeds were preconditioned as described previously (17). Seeds were surface sterilized in 10% (v/v) commercial bleach with agitation, rinsed three times with distilled water, then placed on sterile plates. Sterile distilled water (5 mL) was added, and seeds incubated at 30 °C for 14 days to allow preconditioning.

Preconditioned seeds were then treated with SLA1 at final concentrations of 0.5, 1, 5, 10, and 50 nM. GR24 (Chiralix B.V., Nijmegen, The Netherlands) was included as a positive control. Treated seeds were incubated overnight at 30°C to allow germination. Afterwards, images of *Striga* seedlings were taken using a Leica MZ10F stereomicroscope (Leica Microsystems, Germany) fitted with a DFC320FX camera (Leica Microsystems, Germany) for analysis.

Germination efficiency was calculated as the mean percentage of germinated seeds across three independent fields of view using the formula below:

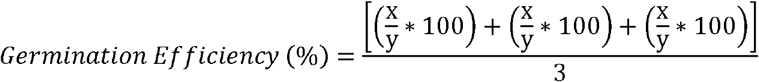

Where x is the number of germinated *Striga* seeds and y is the total number of *Striga* seeds in a specific field of view and 3 is the number of replications.

**Movie S1**. GR24 docked in the ShHTL7 ligand-binding pocket.

**Movie S2**. SLA1 docked in the ShHTL7 ligand-binding pocket.

**Movie S3**. 100-ns molecular dynamics simulation of GR24–ShHTL7 complex.

**Movie S4**. 100-ns molecular dynamics simulation of SLA1–ShHTL7 complex.

**Movie S5**. 100-ns molecular dynamics simulation of GR24–ShHTL9 complex.

**Movie S6**. 100-ns molecular dynamics simulation of SLA1–ShHTL9 complex.

